# Distributional Transformation improves Decoding Accuracy when Predicting Chronological Age from Structural MRI

**DOI:** 10.1101/2020.09.11.293811

**Authors:** Joram Soch

## Abstract

When predicting a certain subject-level variable (e.g. age in years) from measured biological data (e.g. structural MRI scans), the decoding algorithm does not always preserve the distribution of the variable to predict. In such a situation, distributional transformation (DT), i.e. mapping the predicted values to the variable’s distribution in the training data, might improve decoding accuracy. Here, we tested the potential of DT within the 2019 Predictive Analytics Competition (PAC) which aimed at predicting chronological age of adult human subjects from structural MRI data. In a low-dimensional setting, i.e. with less features than observations, we applied multiple linear regression, support vector regression and deep neural networks for out-of-sample prediction of subject age. We found that (i) when the number of features is low, no method outperforms linear regression; and (ii) except when using deep regression, distributional transformation increases decoding performance, reducing the mean absolute error (MAE) by about half a year. We conclude that DT can be advantageous when predicting variables that are non-controlled, but have an underlying distribution in healthy or diseased populations.

## 1 INTRODUCTION

In recent years, probabilistic modelling (Stephan and Mathys, 2014) and machine learning (Rutledge et al., 2019) have been increasingly applied to psychiatric populations and problems, leading to the creation of a whole new field of research called “Computational Psychiatry” (Huys et al., 2016).

The prediction of human age from biological data holds a large promise for computational psychiatry, because biologically predicted age may serve as an important biomarker for a number of human phenotypes (Cole et al., 2017). For example, brain-predicted age may be an indicator for the memory decline associated with neurodegenerative disorders such as Alzheimer’s disease (AD; Lin et al., 2018).

The 2019 Predictive Analytics Competition^1^ (PAC), held before the the 25th Annual Meeting of the Organization for Human Brain Mapping^2^ (OHBM), addressed this research question by asking teams to predict chronological age of human subjects from raw or preprocessed structural magnetic resonance imaging (sMRI) data using a self-chosen machine learning (ML) approach.

Because brain structure changes significantly when becoming older, age can be predicted from sMRI with considerable precision (Cole et al., 2017), usually quantified as mean absolute error (MAE). Brain-predicted age difference (BPAD), i.e. the difference between age predicted from sMRI and actual age, can either be a sign of “accelerated” (BPAD > 0) or “decelerated” (BPAD < 0) brain aging. Accelerated brain aging has been associated with lower levels of education and physical exercise (Steffener et al., 2016), less meditation (Luders et al., 2016) and an increased mortality risk, among others (Cole et al., 2018).

While early attempts at predicting human age from functional MRI (Dosenbach et al., 2010) have used decoding algorithms such as support vector machines (SVM), more recent ML-based decoding from structural MRI have focused on deep learning (Plis et al., 2014), specifically using convolutional neural networks (CNN) to predict chronological age (Cole et al., 2017, 2018; Jiang et al., 2020), AD disease state (Lin et al., 2018) or even body mass index (Vakli et al., 2020) from anatomical brain data.

With a complex series of linear and non-linear optimizations involved in those decoding algorithms, it is clear that the distribution of predicted values of the target variable (e.g. chronological age) will not be exactly identical to the distribution of those values learned from (i.e. the training data). Here, we introduce distributional transformation (DT), a post-processing method for ML-based decoding, which allows to circumvent this problem by matching predictions to the training distribution.

Applied to out-of-sample ML prediction, DT operates by transforming the distribution of predicted values into the distribution of learned values of the variable of interest. In this way, prediction of the target variable (here: chronological age) is not only achieved by reconstructing it from the test set features (here: structural MRI), but additionally aided by looking at the training set samples, such that predictions are more likely to be in a realistic range for that particular target variable.

In this study, we apply DT to PAC 2019 data, while predicting chronological age using either multiple linear regression (GLM), support vector regression (SVR) and deep neural networks (DNN). In summary, we find that (i) multiple linear regression outperforms all other methods in a low-dimensional feature space and (ii) distributional transformation reduces prediction error for linear regression and SVR, but not DNN regression.

## 2 METHODS

### 2.1 Structural MRI data

Data supplied within the 2019 Predictive Analysis Competition (PAC) included structural scans from *n*_1_ = 2640 training set and *n*_2_ = 660 validation set subjects which were all healthy adults. The analyses reported here exclusively used the pre-processed grey matter (GM) and white matter (WM) density images supplied during the competition (for pre-processing details, see Cole et al., 2017). Covariates included subjects’ gender and site of image acquisition; data were acquired at 17 different sites. Subjects’ age in years was supplied for the training set (2640 values), but not shared and only after the competition released for the validation set (660 values). The ratio of training to validation set size is 4:1 (see Table 1).

**Table 1.** Data dimensions and cross-validation. During the competition (second column), the model was developed using 10-fold cross-validation within the training data, before performance was reported on the withheld data set. In the context of this paper (first column), age is predicted out-of-sample in the validation data without cross-validation.

| out-of-sample prediction | k-fold cross-validation | number of subjects |
| --- | --- | --- |
| training | training | 2376 |
|  | testing | 264 |
| validation | reporting | 660 |

### 2.2 Feature extraction

The Automated Anatomical Labeling (AAL; Tzourio-Mazoyer et al., 2002) atlas parcellates the human brain into 90 cortical and 26 cerebellar regions. We used the AAL label image (supplied with MRIcroN^3^ and also available from the TellMe package^4^) and resliced it to the first pre-processed GM image in order to match image dimensions and voxel size. We then extracted average GM and WM density from all 116 regions from the pre-processed structural images for each subject.

Acquisition site information was transformed into 17 indicator regressors and subject gender information was transformed into a +1/−1 regressor. Together with the extracted GM and WM densities, this constituted design matrices for training and validation data having *p* = 2 × 116 + 17 + 1 = 250 columns (see Figure 1).

**Figure 1.**
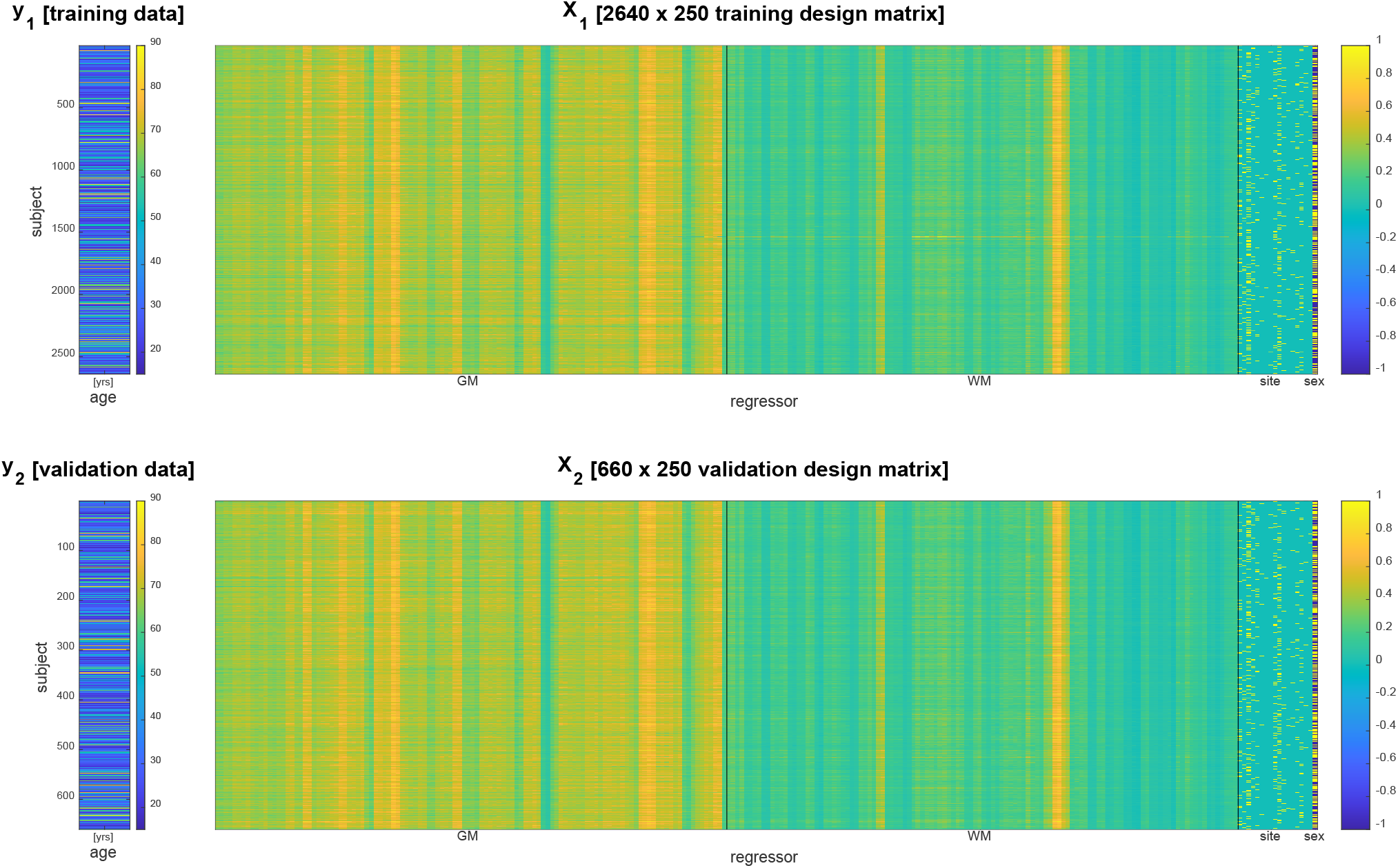
Data vector (left) and design matrix (right) for training set (top) and validation set (bottom). In the design matrices, rows correspond to subjects and columns correspond to brain regions and covariates. Regressors specifying grey matter (GM) and white matter (WM) densities as well as site covariates are separated by vertical black lines.

### 2.3 Decoding algorithms

Let *y*_1_ and *y*_2_ be the *n*_1_ × 1 and *n*_2_ × 1 training and validation data vector and let *X*_1_ and *X*_2_ be the *n*_1_ × *p* and *n*_2_ × *p* training and validation design matrix.

#### 2.3.1 Multiple linear regression

Multiple linear regression proceeds by estimating regression coefficients via ordinary least squares (OLS) from the training data

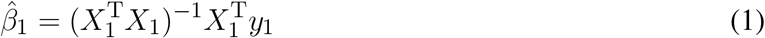

and generating predictions by multiplying the design matrix with estimated regression coefficients in the validation data

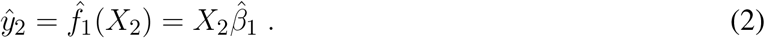

#### 2.3.2 Support vector regression

Support vector regression (SVR) was implemented in MATLAB using fitrsvm. A support vector machine was calibrated using the training data and then used to predict age in the validation data:

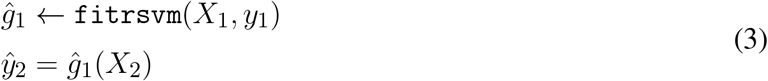

#### 2.3.3 Deep neural network regression

Deep neural network (DNN) regression was implemented in MATLAB using trainNetwork. Before training, non-indicator regressors in *X*_1_ and *X*_2_ were z-scored, i.e. mean-subtracted and divided by standard deviation:

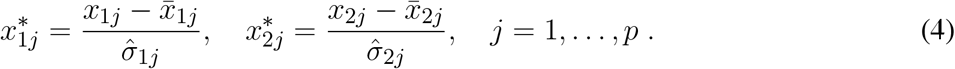

The network consisted of six layers (see Table 2) following a MathWorks tutorial^5^ on deep learning for linear regression and was solved in training using the Adam optimizer. The number of epochs was set to 100, with a mini batch size of 20, an initial learning rate of 0.01 and a gradient threshold of 1. Similarly to SVR, training and prediction proceeded as follows:

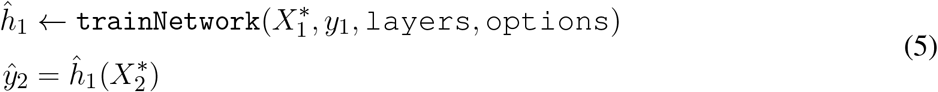

**Table 2.** Layers of the deep neural network. The network employed for DNN regression consisted of six layers which were designated for using deep learning on regression problems.

| MATLAB command | Description | Parameters |
| --- | --- | --- |
| sequenceInputLayer | inputs 1D data into network | 250 features |
| lstmLayer | long short-term memory (LSTM) layer;<br>learns long-range dependencies between features | 125 hidden units |
| fullyConnectedLayer | multiplies with weight matrix and adds bias | 50 output units |
| dropoutLayer | sets elements to zero with given probability | $p = 0.5$ |
| fullyConnectedLayer | multiplies with weight matrix and adds bias | 1 output unit |
| regressionLayer | outputs scalar prediction from network | — |

### 2.4 Distributional transformation

Because the distribution of predicted age values will not exactly match the distribution of validation set age and likely also deviates from the distribution of training set age, one can apply an additional distributional transformation (DT) step after prediction.

DT uses cumulative distribution functions^6^ (CDFs). Let *X* and *Y* be two random variables. Then, *X* is distributionally transformed to *Y* by replacing each observation of *X* by that value of *Y* which corresponds to the same quantile as the original value, i.e.

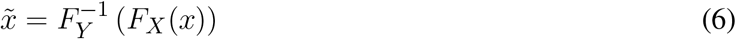

where *F*_*X*_ is the CDF of *X* and 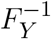 is the inverse CDF of *Y*. Note that DT preserves the complete ordering of *X*, but changes its CDF to that of *Y* (see Appendix A).

Here, we apply DT to the predicted ages *ŷ*_2_, with the goal of mapping them to the distribution of the training ages *y*_1_ by calculating

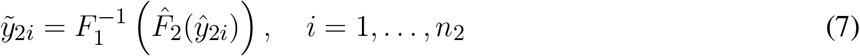

where 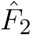 is the empirical CDF of *ŷ*_2_ and 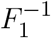 is the inverse empirical CDF of *y*_1_, obtained in MATLAB using `ecdf` (see Appendix B).

After the transformation, the ranks of all predictions *ŷ*_2*i*_ are still the same, but the empirical CDF of 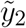 matches that of *y*_1_. In other words, we receive something that looks like the training age values in terms of age distribution, but is still predicted from the validation brain data.

The rationale behind this is that, if training and validation set are unbiased, representative and unsystematic samples from the underlying population, then sampling from the training data should in itself be a good prediction strategy for the validation data (Brodersen et al., 2010). For example, because it can be suspected that mean age has been controlled for when dividing into training and validation data, the age distributions in training and validation data should be close to each other.

### 2.5 Performance assessment

After generating predictions for the validation set, we assessed decoding accuracy using multiple measures of correlation (see Table 3) between predicted ages *ŷ*_2_ and actual ages *y*_2_. During the PAC 2019, Objective 1 was to minimize the mean absolute error. Objective 2 was to minimize Spearman’s rank correlation coefficient between *y*_2_ and (*y*_2_ − *ŷ*_2_), as it is desirable that the brain-predicted age difference (BPAD) is not correlated with age. After assessing performance in the validation set, we conducted several statistical analyses.

**Table 3.** Measures of prediction performance. For PAC 2019 Objectives, see main text.

| Measure | Description |
| --- | --- |
| $R^2$ | coefficient of determination ("R-squared") |
| $R^2_{\text{adj}}$ | adjusted coefficient of determination |
| $r$ | Pearson correlation coefficient |
| $r_{\text{SC}}$ | Spearman's rank correlation coefficient |
| MAE | mean absolute error (Objective 1) |
| RMSE | root mean squared error |
| Obj. 2 | Objective 2 from PAC 2019 |

### 2.6 Statistical analyses

First, we submitted absolute errors (AE) between actual and predicted age to Wilcoxon signed-rank tests^7^ in order to test for significant reduction of the MAE between decoding algorithms (linear regression, SVR, DNN) and prediction methods (with and without DT). This non-parametric test was chosen due to the presumably non-normal distribution of absolute errors.

Second, we calculated the empirical Kullback-Leibler (KL) divergence^8^ of the distribution of actual ages from the distributions of predicted ages. The KL divergence is a non-negative distance measure for probability distributions; the more similar two distributions are, the closer it is to zero. Thus, we expect substantial reductions of the KL divergence after applying DT.

Third, we ran two-sample Kolmogorov–Smirnov (KS) tests^9^ between predicted age values and validation set ages, against the null hypothesis that predicted values and validation ages are from the same continuous distribution. Consequently, similarly to the KL divergence, we expect less significant or non-significant results from the KS test after applying DT.

Finally, for purely illustrative purposes, we investigated the influence of model parameters for the most successful method of age prediction (linear regression with DT). To this end, we concatenated training and validation data (because no out-of-sample testing was needed for this analysis) and calculated parameter estimates for regression coefficients and noise variance

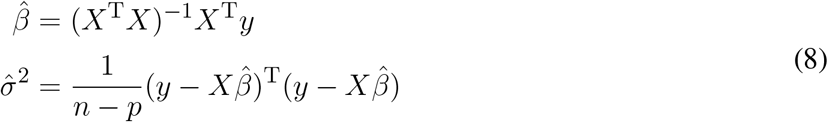

which were then used to calculate standard error and confidence interval for each estimated model parameter (Ashburner et al., 2003, ch. 7, eq. 42; ch. 8, eq. 9)

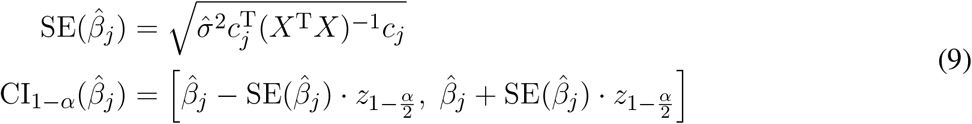

where *c*_*j*_ is a contrast vector of only zeros except for a single one in *j*-th position (i.e. testing *β*_*j*_ against 0), *z*_1−*p*_ is the (1 *− p*)-quantile from the standard normal distribution (”z-score”) and the confidence level was set to (1 *− α*) = 90% (such that 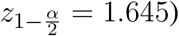.

The complete data analysis as well as resulting decoding accuracies can be reproduced using the contents of a GitHub repository (see Data Availability Statement).

## 3 RESULTS

### 3.1 Influence of decoding algorithm

Qualitatively, prediction performance can be assessed via scatterplots of predicted age against actual age in the validation set (see Figure 2). Quantitatively, the ranking is similar across all measures of correlation (see Figure 3): multiple linear regression performs best (*r* = 0.91, MAE = 5.07 yrs), but only mildly outperforms deep neural network regression (*r* = 0.89, MAE = 5.18 yrs) and strongly outperforms support vector regression (*r* = 0.83, MAE = 6.82 yrs). This is true for measures which are to be maximized (*R*^2^, 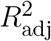, *r*, *r*_SC_) as well as for measures which are to be minimized (MAE, RMSE, Obj. 2). Wilcoxon signed-rank tests indicated significantly lower absolute errors for linear regression, except when compared to DNN without applying DT (see Table 4).

**Figure 2.**
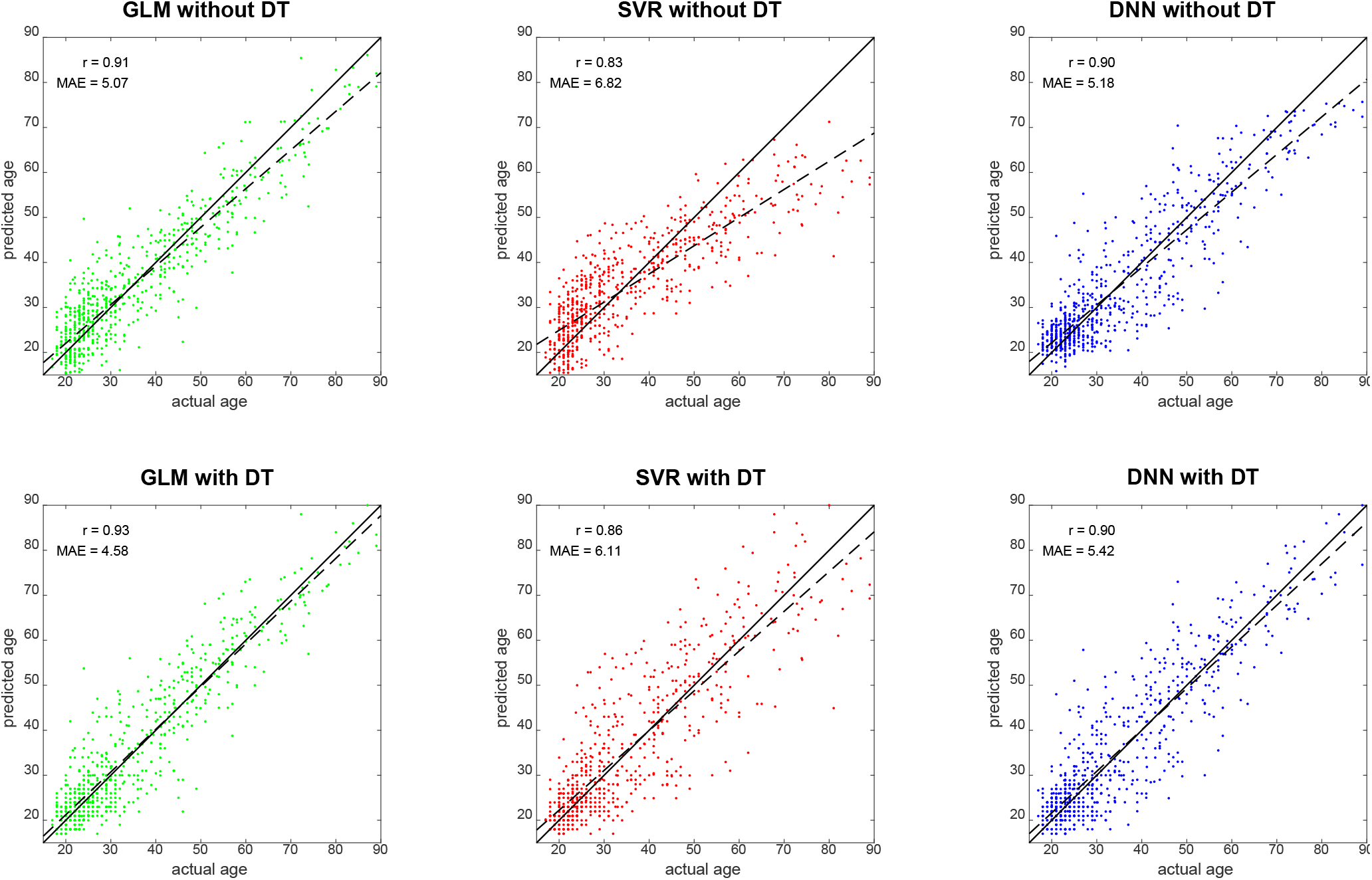
Predicted vs. actual age for validation set subjects. Solid black lines denote the identity function and dashed black lines are regression lines. Also reported are correlation coefficients (*r*) and mean absolute errors (MAE) for each analysis (see Figure 3). Abbreviations: GLM = multiple linear regression; SVR = support vector regression; DNN = deep neural network regression; DT = distributional transformation.

**Figure 3.**
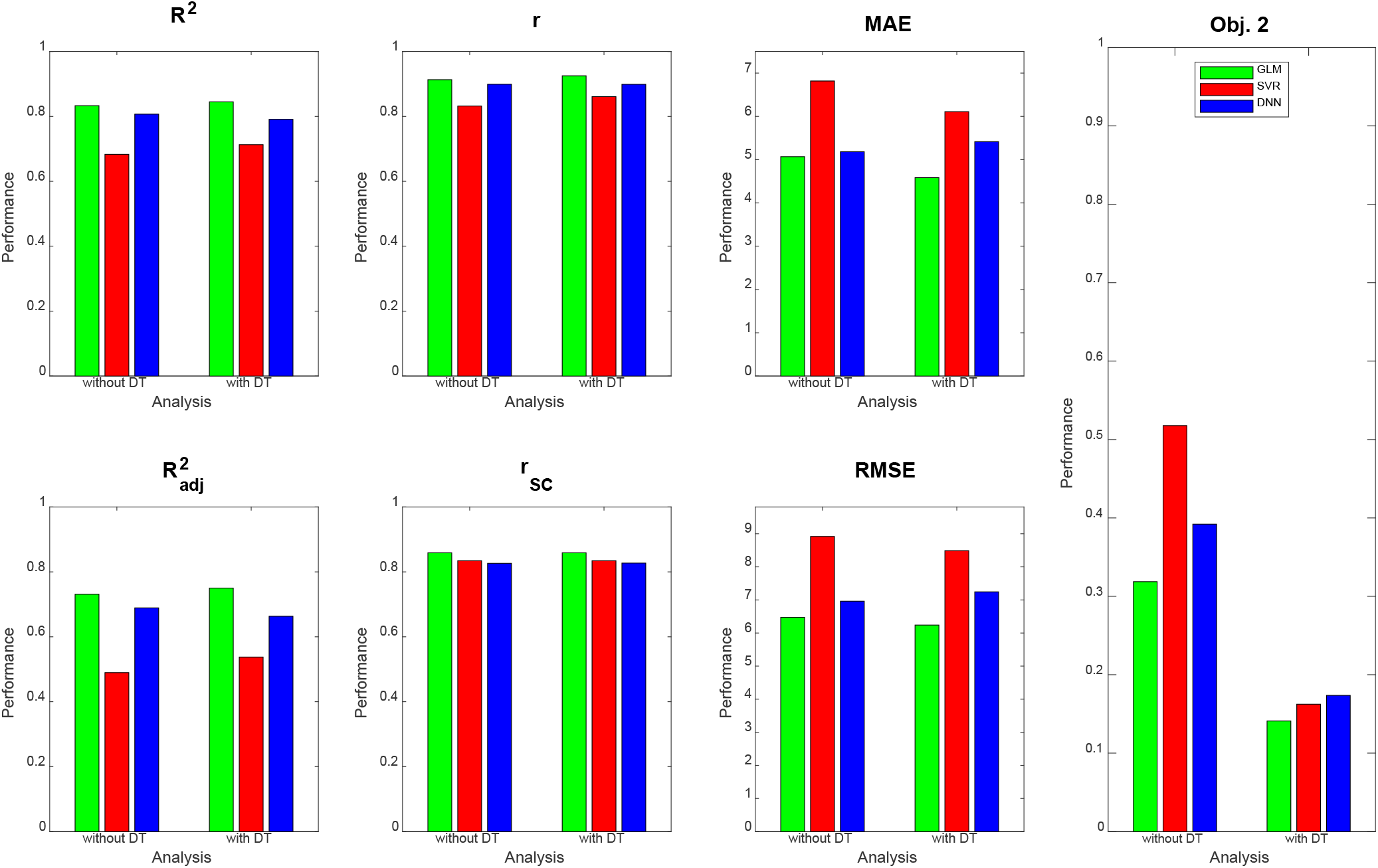
Prediction performance across models, methods and measures. For each performance measure, decoding accuracy is given for multiple linear regression (green), support vector regression (red) and deep neural network regression (blue), with and without distributional transformation (DT). Abbreviations: see Table 3.

**Table 4.** Prediction performance across models and methods. The first row lists performance measures for three decoding algorithms (GLM, SVR, DNN) and two prediction methods (without DT, with DT). The second row reports results from Wilcoxon signed-rank tests comparing GLM against SVR and DNN (thus no entries in the GLM column). The third row reports results from Wilcoxon signed-rank tests comparing each decoding algorithm with and without DT. Negative *z*-values indicate significantly lower absolute errors for GLM (second row) or DT (third row), respectively. Abbreviations: GLM = multiple linear regression; SVR = support vector regression; DNN = deep neural network regression; DT = distributional transformation.

| Statistic | Method | GLM | SVR | DNN |
| --- | --- | --- | --- | --- |
| prediction performance | without DT | $r = 0.91, \text{MAE} = 5.07$ | $r = 0.83, \text{MAE} = 6.82$ | $r = 0.90, \text{MAE} = 5.18$ |
| | with DT | $r = 0.93, \text{MAE} = 4.58$ | $r = 0.86, \text{MAE} = 6.11$ | $r = 0.90, \text{MAE} = 5.42$ |
| comparison of models | without DT | – | $z = -7.44, p < 0.001$ | $z = 0.23, p = 0.817$ |
| | with DT | – | $z = -5.95, p < 0.001$ | $z = -4.12, p < 0.001$ |
| comparison of methods | with DT vs. without DT | $z = -4.90, p < 0.001$ | $z = -4.78, p < 0.001$ | $z = 4.04, p < 0.001$ |

### 3.2 Influence of distributional transformation

When comparing predicted against actual age, one can see that SVR and DNN predictions deviate quite some amount from the actual distribution (see Figure 2, top-middle and top-right), especially by not predicting very high ages (SVR: max(*ŷ*_2_) = 71.25 yrs; DNN: max(*ŷ*_2_) = 75.68 yrs), whereas linear regression creates a more homogeneous picture (see Figure 2, top-left), but also predicts very low ages (min(*ŷ*_2_) = 1.06 yrs).

DT improves the decoding accuracy of linear regression (MAE: 5.07 → 4.58 yrs) and SVR (MAE: 6.82 → 6.11 yrs), reducing their MAE by about half a year. For DNN, the error actually goes up (MAE: 5.18 → 5.42 yrs). Similar results are observed when considering other measures. Wilcoxon signed-rank tests indicated significantly lower absolute errors when applying DT after linear regression and SVR and significantly higher absolute errors when applying DT after DNN regression (see Table 4).

DT especially benefits Objective 2 of PAC 2019, as the Spearman correlation of brain-predicted age difference with age itself goes down considerably for all decoding algorithms when applying DT (see Figure 3, right), thus increasing the independence of prediction error from predicted variable.

When not applying DT (see Figure 4, top row), DNN yields the smallest KL divergence (KL = 0.073), with predictions almost being in the correct range (17 ≤ *y*_2_ ≤ 89), whereas linear regression achieves lower correspondence and SVR suffers from making a lot medium-age predictions (40 ≤ *ŷ*_2_ ≤ 60) and there not being a lot middle-aged subjects in the training and validation sample. When applying DT (see Figure 4, middle row), all methods give rise to the same histogram of predicted ages and have the same and minimally possible distance to the actual distribution (KL = 0.027). Still, despite having the same distribution, prediction performance differs between methods (see Figure 3).

**Figure 4.**
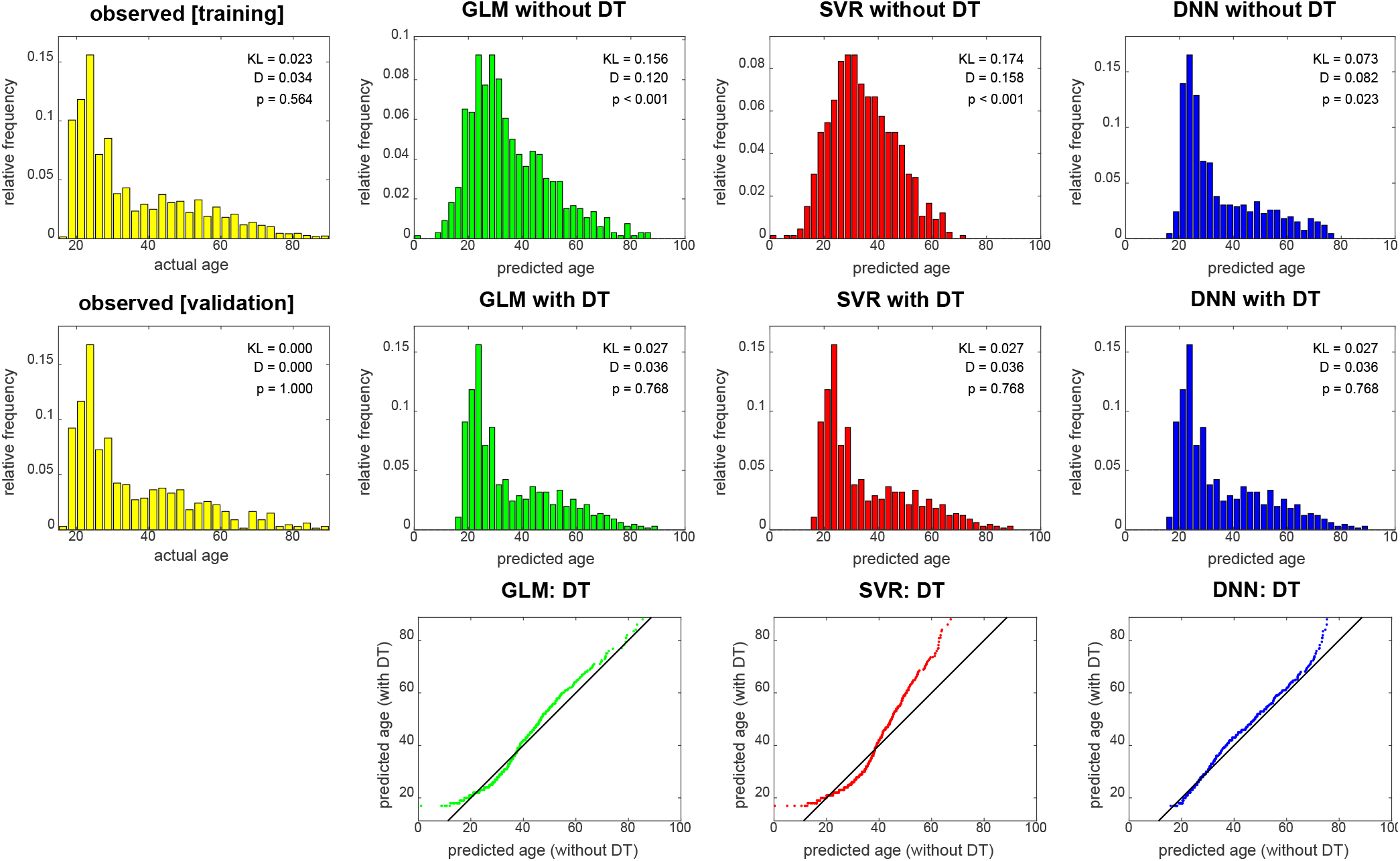
Empirical distributions of actual age (yellow), predicted age (top row) and transformed age (middle row). Each panel shows relative frequencies, i.e. number of occurences divided by total number of subjects. The applied distributional transformation is shown for each decoding algorithm (bottom row), with the diagonal line denoting the identity function. The empirical KL divergence was always computed relative to the validation age distribution (middle row, left), thus this distribution has KL = 0. Also reported are statistics from a two-sample Kolmogorov-Smirnov test between predicted values and validation ages, thus *D* = 0 and *p* = 1 for validation set age. Abbreviations: GLM = multiple linear regression; SVR = support vector regression; DNN = deep neural network regression; DT = distributional transformation; KL = Kullback-Leibler divergence.

These findings are also reflected in results from KS tests which indicate significant differences of actual and predicted age distributions before (see Figure 4, top row), but not after (see Figure 4, middle row) DT was applied to predicted age values. Moreover, it can be seen in the graphical display of the distributional transformations themselves (see Figure 4, bottom row) which deviate stronger from the diagonal line for higher KL divergences before application of DT (see Figure 4).

### 3.3 Influence of regression coefficients

In order to see which features aided successful prediction of age when using multiple linear regression, we report parameter estimates and compute confidence intervals (see Figure 5). These results show that (i) there was no effect of gender on age, putatively because gender was controlled when splitting the data; (ii) there were only mild site effects, putatively because the whole age range was sampled at each site; and (iii) regional GM and WM densities both contributed to the predictions, as variables from both groups have significant effects on subject age (see Figure 5).

**Figure 5.**
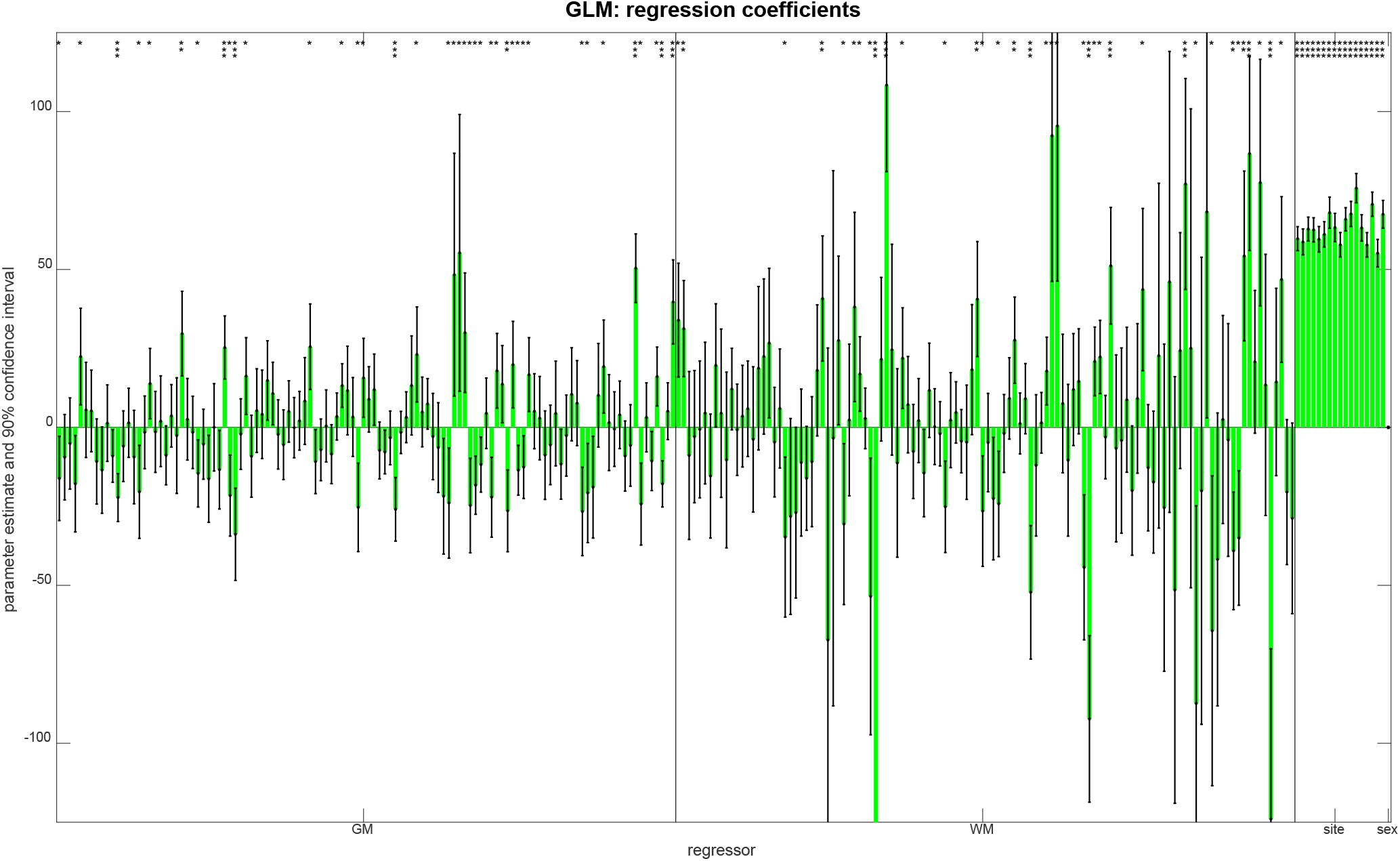
Parameter estimates and confidence intervals from linear regression, obtained when combining training and validation data into one data set. For each regressor, the estimated coefficient is plotted as a bar and a 90 % confidence interval is reported as well. Model parameters are also reported as being significantly different from zero (* *p* < 0.05; ** *p* < 0.001; *** *p* < 0.05/250).

## 4 DISCUSSION

We have applied distributional transformation (DT), a post-processing method for prediction analyses based on machine learning (ML), to predict chronological age from structural MRI scans in a very large sample of healthy adults (Cole et al., 2017). By using DT, we were able to significantly reduce the mean absolute error (MAE) of linear regression and support vector regression, but not deep learning regression.

### 4.1 Distributional transformation

DT can be particularly advantageous when predicting variables which are hard to control experimentally (esp. biological phenotypes), but the distribution of which is known through the availability of training data. The rationale behind distributional transformation for ML-based prediction is simple:

1. A lot of target variables have a natural range into which their values must fall:

a. Human age cannot be smaller than zero (or at least, smaller than 9 months), is rarely larger than 100 years and has thus far not exceeded 122 years.
b. Intelligence quotients (IQ) are (by construction) normally distributed with mean 100 and standard deviation 15.
c. Physical parameters such as weight and height fall into typical ranges differing by gender and age.
d. Probabilities and frequencies, e.g. proportions of correct responses, are bound to the interval [0, 1].
2. When associations between target variable and feature space are learned by sending training samples through a complex machinery of linear and non-linear optimizations, some test set predictions will likely be outside these areas, thereby violating the natural range of the target variable.
3. Distributional transformation brings the predictions back into the natural range by putting them in reference to the training set samples, but preserving the ranks obtained when reconstructing from the test set features.

The extent to which this transformation will work naturally depends on how precise the cumulative distribution functions of training samples and predicted values can be estimated. Generally speaking, those estimates will be more precise, the more samples are available to generate them.

Note that DT assumes independent subsets and identical distributions. This means, (i) training and validation set must be independent from each other in order not to introduce dependencies between data used for training an algorithm and data used for reporting its performance; and (ii) training and validation samples must be drawn from the same underlying distribution in order to justify the assumption that they have the same cumulative distribution function. The first requirement is usually met when samples are independent from each other (e.g. subjects); the second requirement is usually met when the variable of interest (e.g. age) does not influence whether samples are in the training or validation set.

With the present data set, we were able to show that DT improves age prediction from structural MRI using some methods (i.e. linear regression or SVR), reducing the MAE by about half year (see Figure 2, left and middle). Notably, DT does not increase prediction precision when the decoding algorithm (e.g. DNN regression) generates test set predictions that already have a similar distribution as the training set samples. This can also be seen from the fact that DT does not substantially change the distribution of DNN predictions (see Figure 4, right) which is in clear contrast to linear regression and SVR.

It is also noteworthy that DT substantially reduced the correlation between brain-predicted age difference (BPAD) and actual age (see Figure 3, right). This is a highly desirable property, because it means that the prediction error is less dependent on subjects’ age and prediction tends to work as good for a 20-year-old as it works for an 80-year-old adult – which is why this quantity was an objective in the PAC 2019 (see Section 2.5) and this finding makes our work complementary to other approaches attempting to reduce bias in brain age estimation (Cole et al., 2018; Beheshti et al., 2019).

Our explanation for the observed reduction is that DT distributes predicted values more evenly across the age spectrum, thereby avoiding negative prediction errors for older subjects (not predicted as being old) and positive prediction errors for younger subjects (not predicted as being young) – a phenomenon commonly observed (see e.g. Beheshti et al., 2019, Fig. 1). This is also compatible with the fact that linear regression covered the age range more broadly, especially for old ages (see Figure 2, left and Figure 4, top), and achieved the lowest Spearman correlation coefficient with and without applying DT.

### 4.2 Limitations

The limitations of our study are three-fold:

- First, we were operating in a low-dimensional setting with fewer features than observations (here: *n* = 2640 > 250 = *p*). Our analyses therefore do *not* show that DT also improves decoding accuracy in high-dimensional settings (where *n < p*) such as decoding from voxel-wise structural data. Previous studies suggest that DNNs outperform simpler methods in this regime (Plis et al., 2014; Cole et al., 2017), but this does not preclude that DT further improves accuracy of CNN predictions.
- Second, we were performing feature extraction using an *a priori* selected brain atlas (here: by extracting from AAL regions). Our analyses therefore do *not* show that DT also improves decoding accuracy under other feature extraction methods or after feature dimensionality reduction. The results reported in this study suggest that DT works well with region-based feature extraction and relatively simple decoding algorithms (linear regression, SVR), but that does not preclude that DT also improves prediction after voxel-based feature extraction.
- Third, we were exclusively analyzing data from healthy subjects (here: by using PAC 2019 data). Our results therefore do *not* apply to clinically relevant groups such as subjects suffering from Alzheimer’s disease (AD) or mild cognitive impairment (MCI). Because structural MRI data contain signatures of chronological age and disease status in patients as well (Lin et al., 2018), we expect DT to also show its merits in those clinical contexts – provided that training and validation set constitute representative samples from the underlying population.

### 4.3 Conclusion

Our results suggest that, when combining distributional transformation with relatively simple decoding algorithms (e.g. linear regression or SVR), predicting chronological age from structural MRI can reach acceptable decoding accuracies in short time. We have provided an algorithm for DT (see Appendix) that can be easily added as a post-processing step to ML-based prediction analyses. Future studies may investigate whether the DT methodology might also be beneficial in other areas of computational psychiatry than brain age prediction.

## 5 APPENDIX

## A Proof of distributional transformation

Let *X* and *Y* be two random variables. Then, *X* is distributionally transformed to *Y* by replacing each observation of *X* by that value of *Y* which corresponds to the same quantile as the original value (see Section 2.4), i.e.

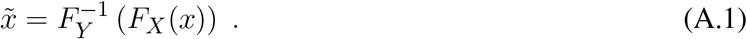

where *F*_*X*_(*x*) is the cumulative distribution function (CDF) of *X* and 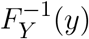is the inverse CDF of *Y*. Consequently, the CDF of 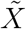 follows as

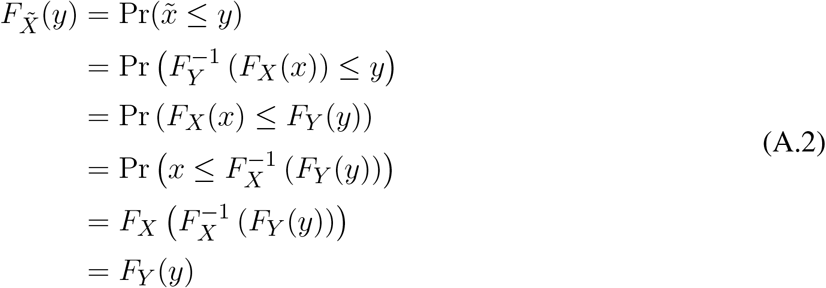

which shows that 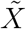 and *Y* have the same CDF and are thus identically distributed.

## B Code for distributional transformation

The following code distributionally transforms x to y in MATLAB:

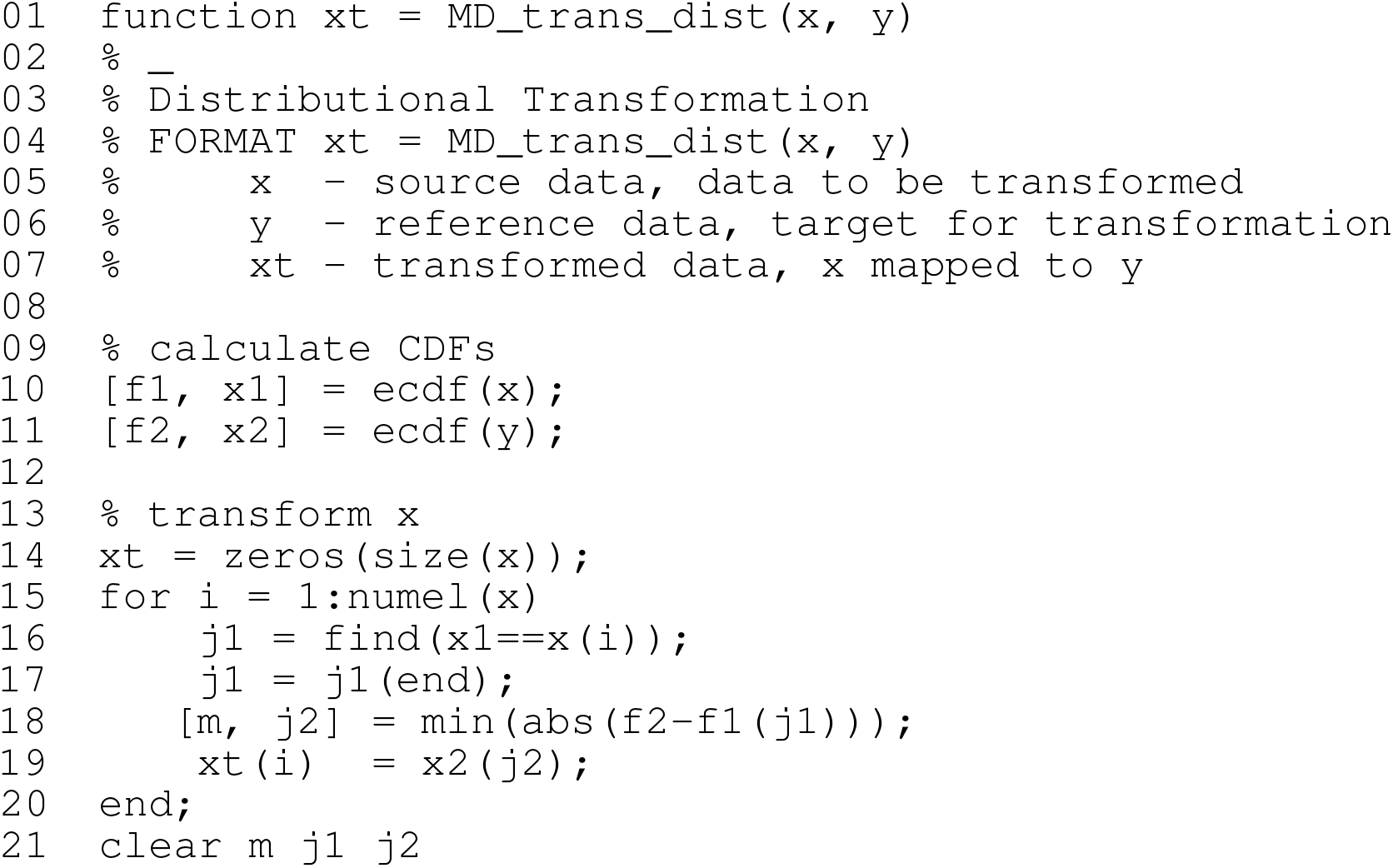

## AUTHOR CONTRIBUTIONS

JS conceived, implemented and performed data analysis; created figures and tables; and wrote the paper.

## ACKNOWLEDGMENTS

The author would like to thank Carsten Allefeld for discussing the linear regression analysis and the distributional transformation method.

We acknowledge support from the German Research Foundation (DFG) and the Open Access Publication Fund of Charité – Universitätsmedizin Berlin.

## DATA AVAILABILITY STATEMENT

Data analyzed in this study were available online during the 2019 Predictive Analytics Competition. For this work, no raw images, but only pre-processed maps were used (see Section 2.1). Requests for data release should be directed to Tim Hahn^10^ and Ramona Leenings^11^.

MATLAB code for (i) feature extraction from pre-processed data, (ii) decoding analyses underlying the results presented in this paper and (iii) results display to reproduce the figures shown in this paper can be found in an accompanying GitHub repository^12^.

## Footnotes

1 https://www.photon-ai.com/pac2019

2 https://github.com/ohbm/OpenScienceRoom2019/issues/10

3 https://people.cas.sc.edu/rorden/mricron/install.html

4 https://github.com/JoramSoch/TellMe

5 https://de.mathworks.com/help/deeplearning/ug/sequence-to-sequence-regression-using-deep-learning.html, accessible in MATLAB as `openExample(’nnet/SequencetoSequenceRegressionUsingDeepLearningExample’)`

6 https://statproofbook.github.io/D/cdf

7 https://de.mathworks.com/help/stats/signrank.html

8 https://statproofbook.github.io/D/kl

9 https://de.mathworks.com/help/stats/kstest2.html

10

11

12 https://github.com/JoramSoch/PAC2019

